# Interspecies Chimeric Conditions Affect the Developmental Rate of Human Pluripotent Stem Cells

**DOI:** 10.1101/2020.09.12.293357

**Authors:** Jared Brown, Christopher Barry, Matthew T. Schmitz, Cara Argus, Jennifer M. Bolin, Michael P. Schwartz, Amy Van Aartsen, John Steill, Scott Swanson, Ron Stewart, James A. Thomson, Christina Kendziorski

**Author notes:** These authors contributed equally to this work (Jared Brown, Christopher Barry). These authors supervised this work equally (James A. Thomson, Christina Kendziorski). Corresponding Authors: Christina Kendziorski, Department of Biostatistics and Medical Informatics, University of Wisconsin-Madison, WI, USA, Christopher Barry, Morgridge Institute for Research, Madison, WI, USA.

## Abstract

Human pluripotent stem cells hold significant promise for regenerative medicine. However, long differentiation protocols and immature characteristics of stem cell-derived cell types remain challenges to the development of many therapeutic applications. In contrast to the slow differentiation of human stem cells *in vitro* that mirrors a nine-month gestation period, mouse stem cells develop according to a much faster three-week gestation timeline. Here, we tested if co-differentiation with mouse pluripotent stem cells could accelerate the differentiation speed of human embryonic stem cells. Following a six-week RNA-sequencing time course of neural differentiation, we identified 929 human genes that were upregulated earlier and 535 genes that exhibited earlier peaked expression profiles in chimeric cell cultures than in human cell cultures alone. Genes with accelerated upregulation were significantly enriched in Gene Ontology terms associated with neurogenesis, neuron differentiation and maturation, and synapse signaling. Moreover, chimeric mixed samples correlated with *in utero* human embryonic samples earlier than human cells alone, and acceleration was dose-dependent on human-mouse co-culture ratios. Differences in the timing and expression levels of genes corresponding to neuron cell types and brain region identity under chimeric conditions were also observed. The altered developmental rates and lineage outcomes described in this report have implications for accelerating human stem cell differentiation and the use of interspecies chimeric embryos in developing human organs for transplantation.

**Author Summary:** Human pluripotent stem cells often require long *in vitro* protocols to form mature cell types of clinical relevance for potential regenerative therapies, a ramification of a nine-month developmental clock *in utero* that also runs *ex utero*. What controls species-specific developmental time and whether the timer is amenable to acceleration is unknown. Further, interspecies chimeric embryos are increasingly being created to study early human development or explore the potential growth of human organs for transplantation. How the conflicting developmental speeds of cells from different species co-differentiating together affect each other is not understood. Here, using genome-wide transcriptional analysis of RNA-sequencing time courses, we show that 1) co-differentiating human embryonic stem cells intermixed with mouse stem cells accelerated elements of human developmental programs, 2) the acceleration was dose-dependent on the proportion of mouse cells, and 3) human cells in chimeric samples correlated to *in utero* samples earlier than human only samples. Our results provide evidence that some components of species-specific developmental clocks may be susceptible to acceleration.

## Introduction

Mammals develop to tremendously different sizes at vastly different rates in the embryo. Little is known about the mechanisms regulating embryonic developmental rates, but they are not uniformly tied to animal size. For example, the smallest mammal, the Etruscan shrew, is approximately one eighth the birthweight of the mouse yet requires 27 rather than 20 days of gestation. The hippopotamus is born a full month before a human infant yet is over ten times heavier, and the largest mammal, the blue whale, has a mass 27 times greater than that of the African elephant at birth despite requiring half the gestational time ^1–3^.

Curiously, when pluripotent stem cells are cultured *in vitro*, they retain the developmental timing of their species of origin despite the lack of maternal factors, suggesting the existence of an intrinsic developmental clock ^4–10^. Currently, the nature of the species-specific developmental clock, including the extent to which it can be warped, is unknown ^11^. The retention of a slow differentiation rate that reflects a nine-month human gestation timeline often results in long differentiation protocols and immature cell characteristics that impede many potential clinical applications of human pluripotent stem cells ^12,13^.

In contrast to the slow differentiation of human stem cells, mouse stem cells differentiate substantially more quickly, reflecting a 20-day rather than a nine-month gestation timeline ^8,14,15^. For example, mature neurons are produced in only 5-14 days from mouse ES cells, while the same cell types can take several months to generate from human embryonic stem (hES) cells ^10,16–18^. Previously, we found that hES cell differentiation was not accelerated in teratomas developed in a mouse despite being exposed to murine host factors ^4^. However, we did not test whether factors active during murine embryonic development could be sufficient to accelerate hES cell differentiation.

Here, we investigated whether hES cells co-differentiated among mouse pluripotent stem cells could accelerate their developmental rate. Under neural differentiation of chimeric co-cultures, we found earlier upregulation and peak expression of hundreds of genes involved in neurogenesis, neuron maturation, and synapse signaling compared to hES cells alone. The accelerated effects were dose-dependent on the starting ratios of human-mouse cells in co-cultures, and chimeric cultures correlated to *in utero* human embryonic samples earlier than human cells alone. We also describe temporal differences in gene expression levels corresponding to brain region identity, suggesting there may be some lineage outcome effects from chimeric co-culture conditions. Overall, we demonstrate that chimeric human-mouse culture conditions are sufficient to accelerate some elements of human stem cell differentiation.

## RESULTS

### Comprehensive RNA-sequencing time course of neural differentiation in chimeric human-mouse co-cultures

We previously described a detailed RNA-sequencing (RNA-seq) time course of mouse and human pluripotent stem cells over three- or six-weeks of neural differentiation, respectively, to characterize the drastically different species-specific rates of development *in vitro* ^4^. Here, we set out to determine if co-differentiating human cells with mouse cells together could induce the human cells to differentiation at a quickened pace. Since hES cells are thought to more closely represent a post-implantation pluripotent stage, we used the similarly-staged mouse Epiblast stem (mEpiS) cells to compare with H9 hES cells ^19–21^. To identify cells from each species, we used mEpiS cells constitutively expressing cytoplasmic efficient green fluorescent protein (EGFP) and H9 cells expressing nuclear-localized H2B-mCherry (Fig. 1).

**Figure 1:**
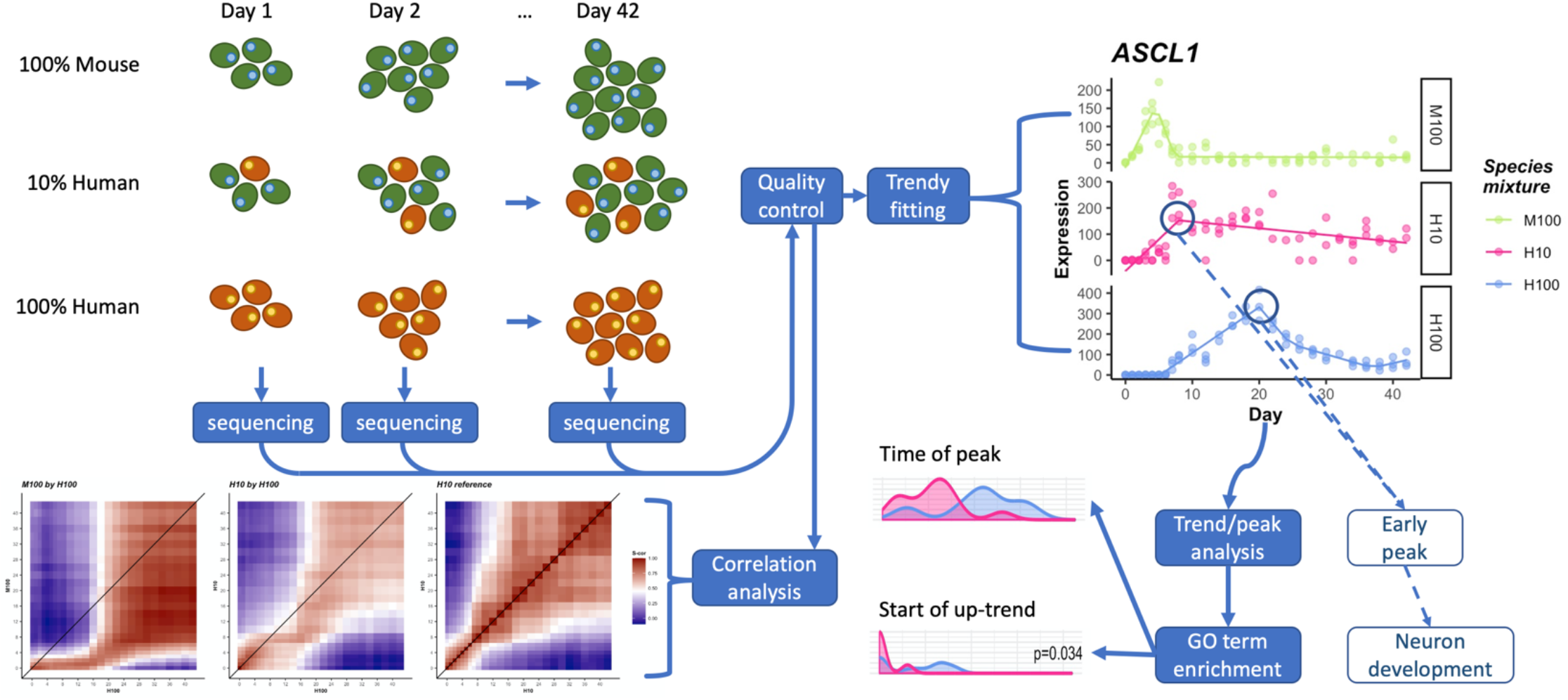
Overview of data collection/analysis pipeline. (top left) Human (red) and mouse (green) cells are cultured in various mixing proportions over the course of 42 days. Every 1-2 days, tissue samples are taken from each time course and sequenced to generate three time-courses of RNA expression data. Low quality biological replicates are removed from analysis and the data are normalized. (top right) Normalized data are fit to segmented regression built for RNAseq data (Trendy) and temporal gene characteristics, such as peak times, are identified. (bottom right) Classified gene sets are passed on for further analysis, in particular, enrichment analysis for GO terms which are temporally accelerated or otherwise systematically altered in H10 compared to H100. (bottom left) In parallel to the previous analysis, normalized data are also correlated between time courses to identify transcriptome-wide effects. Additionally, normalized data are correlated with in-vivo sequencing data from human neural tissue of known age and origin (shown in Fig 7, Supp. Fig 5).

To maximize any potential mouse-induced effects on human differentiation rate, we began by outnumbering human cells with the more quickly differentiating mouse cells in a ten-to-one ratio. 10% human co-cultured cells (H10), along with 100% mouse (M100) or 100% human (H100) control samples, were cultured under identical neural differentiation culture conditions (see Materials and Methods) and samples in triplicate were collected for RNA-seq every 24 or 48 hours for six weeks (Fig. 1). After aligning transcripts to a combined human-mouse transcriptome to derive species-specific expression from the chimeric samples, samples passing quality control parameters (S1 Fig., see Materials and Methods) were processed for correlation analysis, fitted with gene expression patterns using the segmentation regression analysis R-package Trendy ^22^, and the timing of expression pattern changes were compared across samples (Fig. 1).

Although mouse and human cells were singularized before seeding, time lapse microscopy revealed that cells preferentially associated with cells of their own species (Fig. 2, Movie S1). Flow cytometry analysis revealed that although the intended starting cell ratios were seeded, as mouse cells differentiated quickly to become post-mitotic neurons, the still-proliferating human progenitor cells eventually overtook the culture. By day 12 of differentiation ^~^50% of H10 samples were of human composition, and by day 16 over 75% of samples were human cells (S2 Fig.). Although cells tended to associate and proliferate in species-specific clusters, cells from each species did grow alongside each other and interact (Fig. 2, Movie S1).

**Figure 2:**
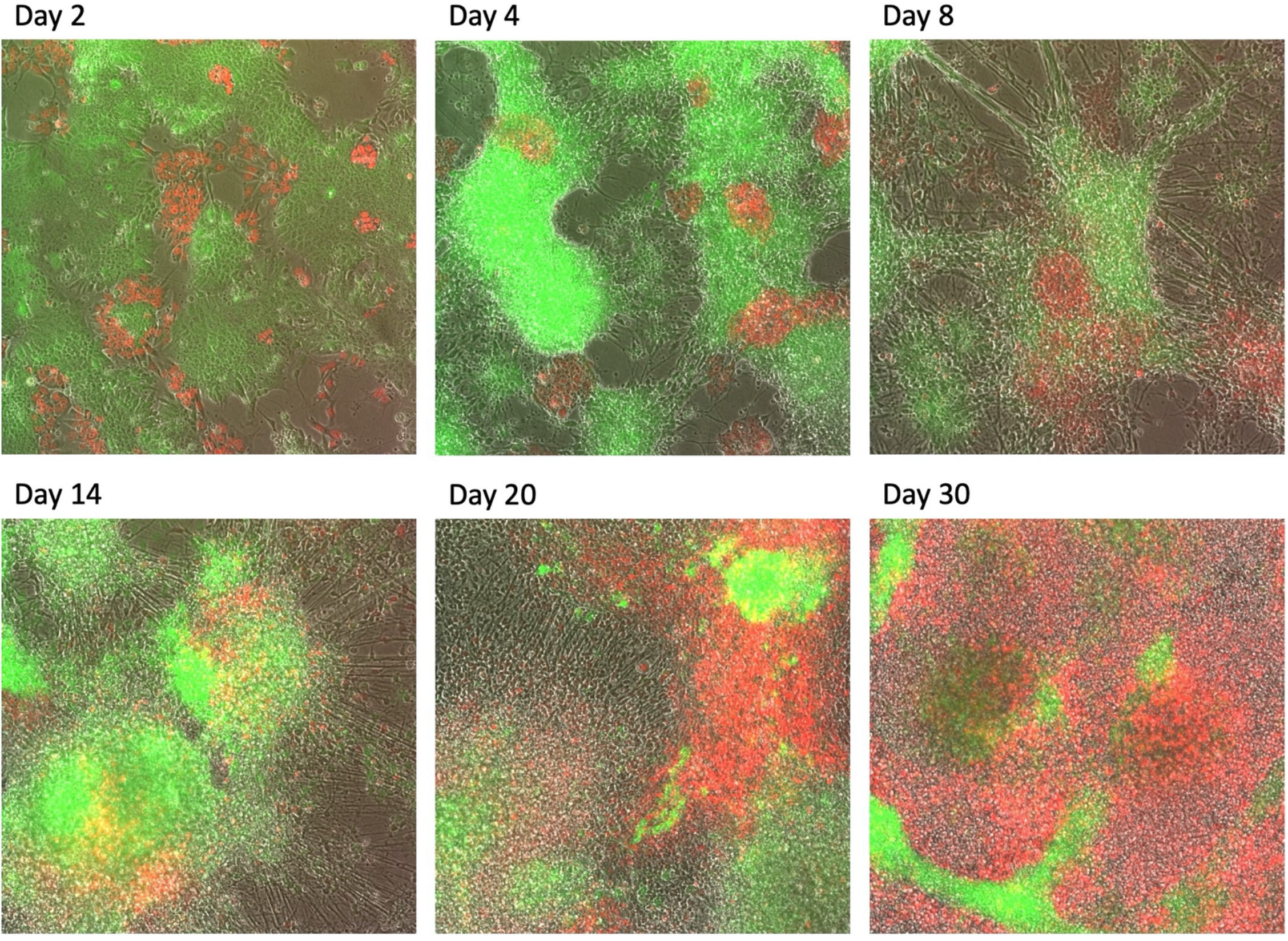
microscopy snapshots of the H10 mixture across the time course. Human cell (H9 cell line) express nuclear-localized H2B-mCherry and so fluoresce red (see methods for details). Mouse cells (cell line derived from C57BL/6-Tg(CAG-EGFP)1Osb/J mice) express EGFP and so fluoresce green (see methods for details). Mouse cells are observed to rapidly differentiate into post-mitotic neurons out to days 8-14. Still differentiating human cells, however, divide at rates after day 8 which quickly sees human cells dominating the human/mouse cell ratio, despite the initial seeding of only 10% human. The simultaneous grouping of human and mouse cells (red and green clusters respectively) suggests preferential association. However, the boundaries between species clusters demonstrate non-zero overlap and interaction.

### Human neurogenic and synaptic genes were upregulated earlier in human-mouse chimeric co-cultures

To determine if gene expression patterns were accelerated in chimeric co-cultures, genes with fitted expression trends were compared between neural differentiation of human cells alone (H100) versus those mixed 1:10 with mouse cells (H10). We first asked if upregulated genes (genes trending up immediately or genes showing no change and then trending up) were upregulated earlier in mixed compared to control samples. Our bioinformatic analysis revealed that 929 genes were upregulated significantly earlier (S1 File) (begin up trending at least 2 days earlier) in H10 versus H100 samples, representing over 41% of all genes that begin as unchanged followed by upregulation (Fig. 3A). We recognized several well-described neurogenic genes identified as accelerated in this early-upregulated category (S3A Fig.), including genes involved in neural differentiation and migration (e.g. STMN2, DCX, NEFL, NEUROG2, MYT1, MAPT), forebrain development (e.g. FEZ1 and EFNB3), neuronal signaling and synapse transmission (e.g. SNAP25, SYT3, SYT4, SYN1), neural stem cell identity (e.g. FABP7, FGF10), and glutamatergic and GABAergic neurons (e.g. SLC1A3, GRIN2D, GABRA1; Fig. 3B). Therefore, genes from a seemingly wide range of neurodevelopmental functions were upregulated earlier under chimeric differentiation conditions.

**Figure 3:**
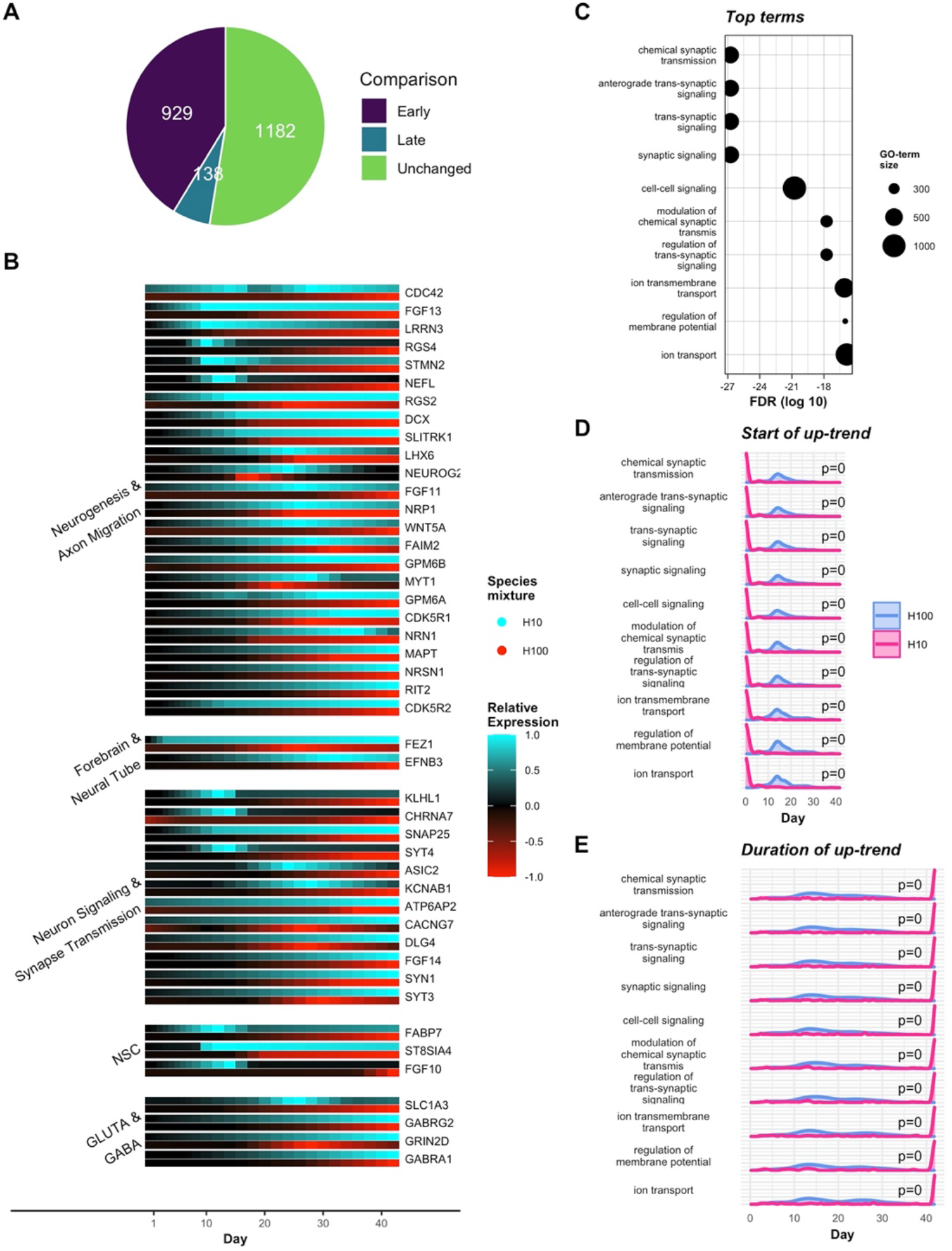
Earlier increases in gene expression in H10 show accelerated activity in signaling gene-sets. (A) All genes which trend up in both H10 and H100 are classified as either early, late, or unchanged in H10 relative to H100. These counts omit genes which start up-regulating between days 0 and 2 in either H10 or H100 as these genes could not be early/late regardless of induced change in expression. (B) Relative expression plots of a curated subset of early-up (EU) genes collected into functional/regional groups. H10 (blue) and H100 (red) time courses are scaled such that 0 expression shows black and maximum expression between H10 and H100 shows as 1/-1 (within gene). (C) Top 10 most significantly enriched GO terms show a strong acceleration in the activation of signaling pathways. Term enrichments are displayed in terms of log10 adjusted p-values (FDR) and sized by the number of genes in the term, not all of which are necessarily classified as EU. (D) Densities of the time of start of the up-trend among (EU) genes in each of the selected GO terms. The distribution of H10 start times (magenta) are tested for significant shift to the left relative to H100 (blue) (KS test). EU genes are observed to typically start increasing from day 0 in the H10 sample. (E) Densities of the duration of the identified up-trend in genes for the same GO terms. Testing for a significant shift to the right in H10 relative to H100 shows the observed pattern to be significant, and EU genes in H10 can be seen to typically increase for the entire time course.

Given that several recognizable neurogenic genes were among those identified as upregulated earlier in H10 compared to H100 samples (Fig. 3B), we set out to statistically test if early upregulated genes were specific to neural differentiation or biasedly identified from a collection of genes within a random assortment of cellular processes. Upon analyzing early-upregulated genes for functional enrichment of GO-terms, we discovered that all of the ten most statistically significantly-enriched terms were associated with neuron and synaptic signaling (Fig. 3C), confirming that neural genes were indeed specifically upregulated earlier in human cells co-differentiated with mouse cells. The beginning of upregulation was not only earlier in these GO-term-associated genes (Fig. 3D), but the duration of up-regulation was also significantly longer, often still trending upwards at the end point of the 6-week time course (Fig. 3E). However, although the up-trend started significantly earlier and lasted longer in chimeric co-cultures, their slopes were also significantly less steep then those of H100 samples (S4A Fig). These results indicate an earlier onset of synaptic signaling gene activation yet a slower rate of upregulation. Taken together, we found that co-culturing human and mouse cells during neural differentiation upregulated genes associated with neuron maturation and synapse formation earlier than human cells alone.

### Regulation of peak gene expression profiles occurs more rapidly in co-cultures with mouse stem cells

During development, genes involved in neural differentiation are often not simply turned on, but rather are expressed in temporally-regulated dynamic patterns ^23,24^. To determine if genes with coordinated expression profiles were regulated more quickly, we next tested whether genes with peak expression profiles (consecutive up-down or up-flat segments) peaked earlier under chimeric versus human control conditions.

Overall, we identified 535 genes that peaked earlier (at least two days) (S1 File) in chimeric culture conditions compared to control samples, representing over 46% of all peaking genes identified in the time course (Fig. 4A). Similarly to early-upregulated genes, we recognized several peaking genes involved in neural development in the accelerated peak category (Fig. 4B and S3B Fig), including genes involved in neurogenesis (e.g. ASCL1, NGFR, NEFM, TUBB3), neural tube development (e.g. MEIS1, GLI3, DLL3), neuron signaling (e.g. SNAP25, ATCAY), and ventral midbrain differentiation (e.g. ISL1, LHX4, NKX6-1). We further validated that genes involved in neurodevelopment were specifically peaking early through GO-term enrichment analysis, and we found that all of the top ten most significantly enriched terms were associated with neural development (Fig. 4C). In contrast to early-upregulated genes which were enriched in neuron and synaptic signaling, early peaked genes were involved in neurogenesis, neuron projection development, and neuron differentiation (Fig. 4C-E). Further, whereas early-upregulated genes had a slower rate of increase compared to control cells, early peaked genes exhibited an earlier time of start of upregulation towards the peak and a faster rate of upregulation to reach the peak (Fig. 4D&E and S4B Fig.). Taken together, genes temporally regulated in peak profiles involved in neurogenesis and neuron differentiation peaked earlier in hES cells co-differentiated with mouse cells than human cells alone, and did so by beginning their upward trend towards the peak earlier and with a steeper slope.

**Figure 4:**
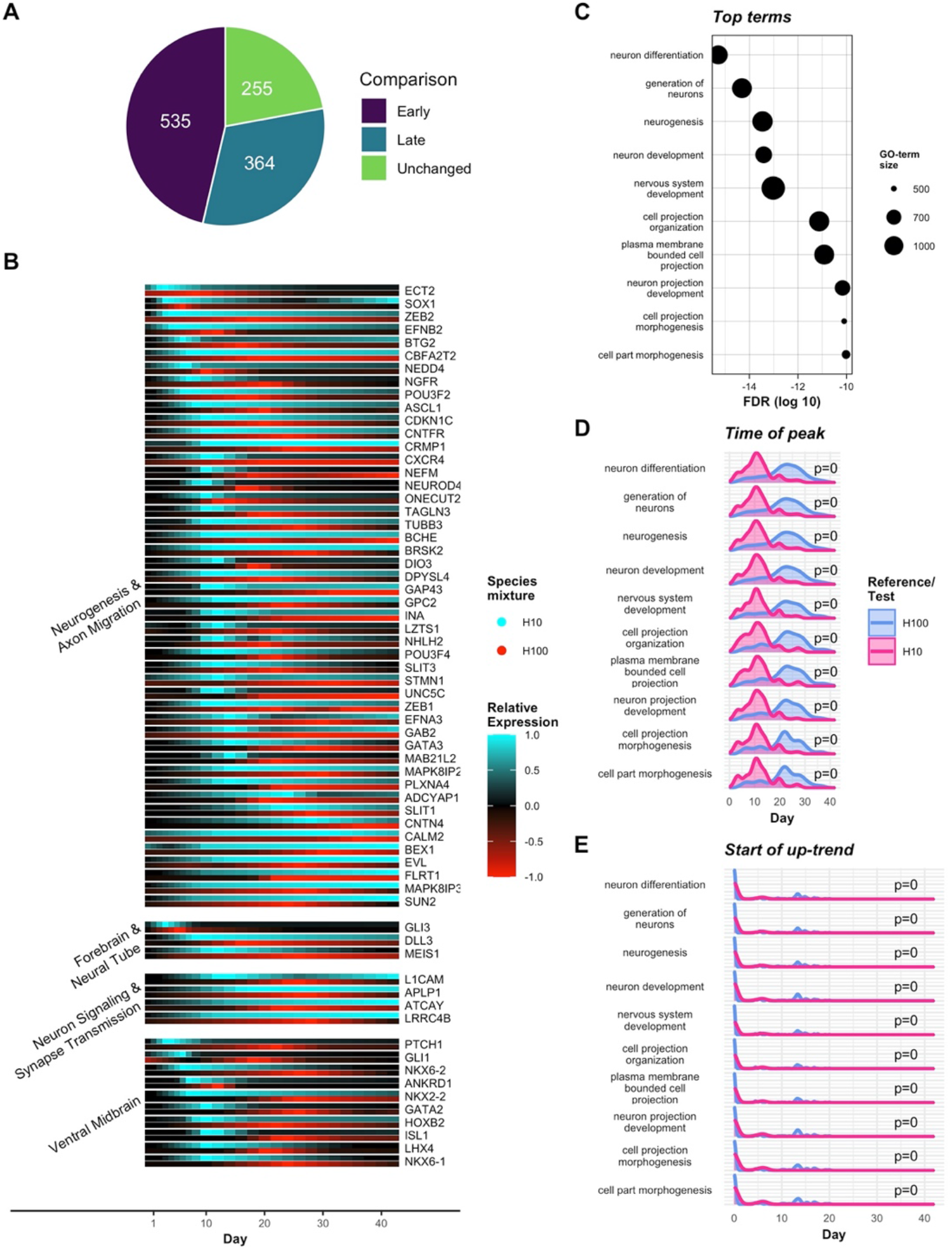
Earlier peaks in gene expression in H10 show accelerated activity in neuron development gene-sets. (A) All genes which peak in both H10 and H100 are classified as either early, late, or unchanged in H10 relative to H100. (B) Relative expression plots of a curated subset of early-peak (EP) genes collected into functional/regional groups. H10 (blue) and H100 (red) time courses are scaled such that 0 expression shows black and maximum expression between H10 and H100 shows as 1/-1 (within gene). (C) Top 10 most significantly enriched GO terms show a strong acceleration in the activation of neuron development pathways. Term enrichments are displayed in terms of log10 adjusted p-values (FDR) and sized by the number of genes in the term, not all of which are necessarily classified as EP. (D) Densities of the time of peak among (EP) genes in each of the selected GO terms. The distribution of H10 start times (magenta) are tested for significant shift to the left relative to H100 (blue) (KS test). (E) Densities of the start of the up-trend leading to the peak in the identified EP genes for the same GO terms. Testing for a significant shift to the left in H10 relative to H100 shows the observed pattern to be significant.

### Chimeric co-culture affected timing and expressions levels of genes associated with neural cell type and brain region identity

Our neural differentiation protocol recapitulates a general neural developmental program and produces neurons of various regional identities ^4^. To determine if chimeric co-culture of hES cells would affect cell lineage outcomes, we identified genes that were most differentially expressed (S1 File) (measured as fold change between maximum expression along the time course) in chimeric mixed samples compared to hES cell controls (Fig. 5).

**Figure 5:**
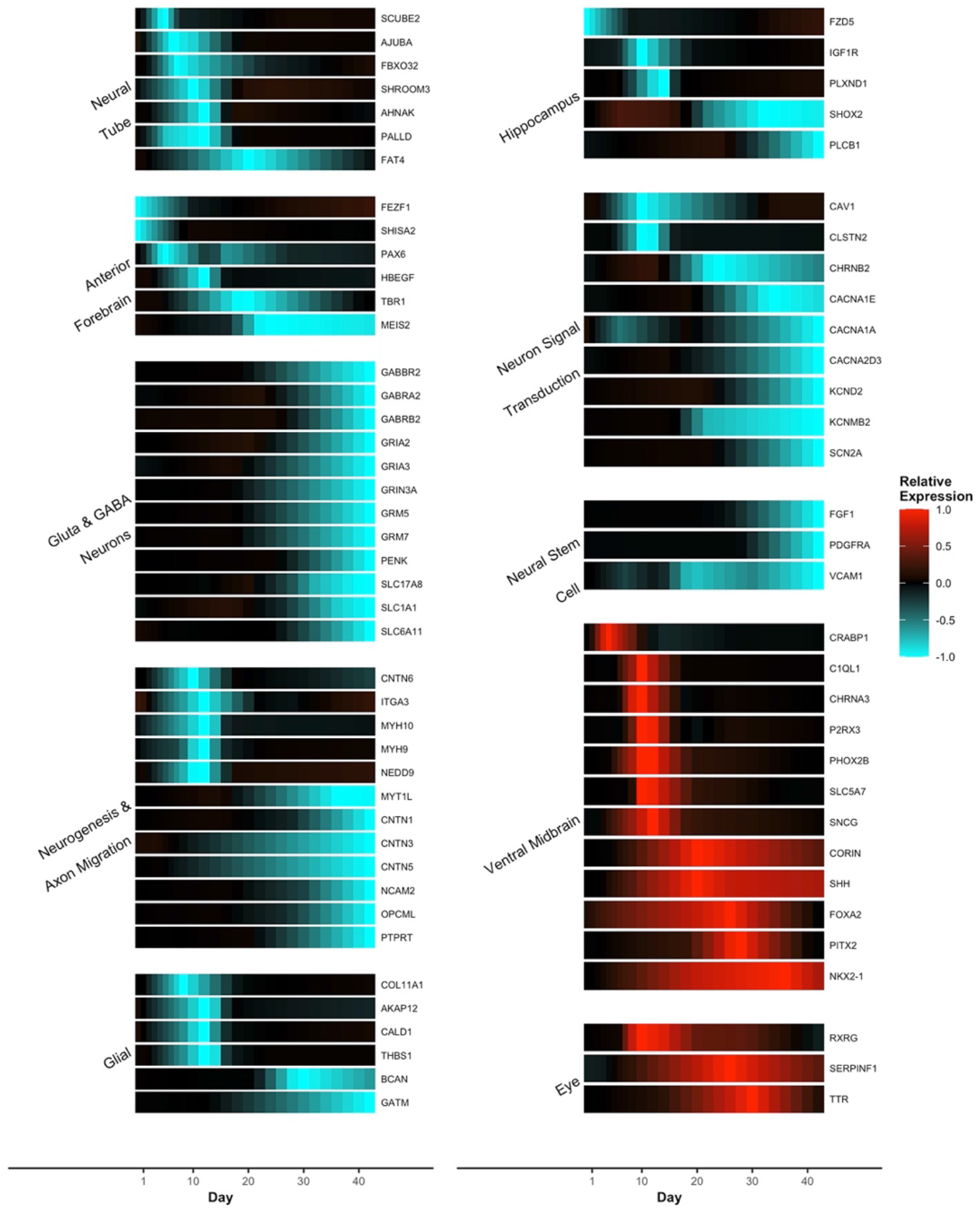
Up/down regulation of genes in H10 show region specific patterns. Relative expressions of curated genes in regional/functional groups are plotted on a normalized −1 to 1 scale. Gene expression (within gene) is normalized such that the maximum difference in fitted expression (in H100 or H10) equals 1. Relative expressions are then calculated as the difference between H10 and H10 where higher H10 values tend towards 1 (red), lower H10 values tend towards −1 (blue), and equivalent values tend towards 0 (black).

We observed some changes in the expression of transient signals as well as changes in sustained region-specific expression. Certain genes associated with the anterior dorsal neural tube and forebrain, glial cells, and the hippocampus showed down regulation in transient periods of gene expression in chimeric conditions compared to human control samples. Similarly, some genes associated with Gluta- and GABAergic neurons and neuron signal transduction showed patterns of sustained downregulation in the later period of the time course (Fig. 5). Other genes associated with neurogenesis and axon migration show a mixture of these patterns. In contrast, some genes associated with the ventral midbrain showed transient upregulation in chimeric mixed samples compared to control samples (Fig. 5). Our analysis therefore revealed that some genes associated with neuron cell type and regional identity were temporally and/or differentially expressed under chimeric conditions.

### Acceleration effects are dose-dependent on percentage of mouse stem cells

Having established significant patterns of altered neuron-associated expression between chimeric mixed samples and control samples, we decided to test the dependence of these results on the initial mixing proportion of human and mouse cells. To this end, we co-cultured human and mouse stem cells in an initial mixing proportion of 85% human and 15% mouse cells (H85) rather than 10% human and 90% mouse cells (H10) to determine if the acceleration effect was dose-dependent (Fig 6A). Sequencing data was collected in triplicate on the same schedule as for the H10, H100, and M100 samples. Identical quality control filtering and segmented regression (Trendy) on the H85 time course produced a dataset directly comparable to the previous mixture/control samples (S2 File).

**Figure 6:**
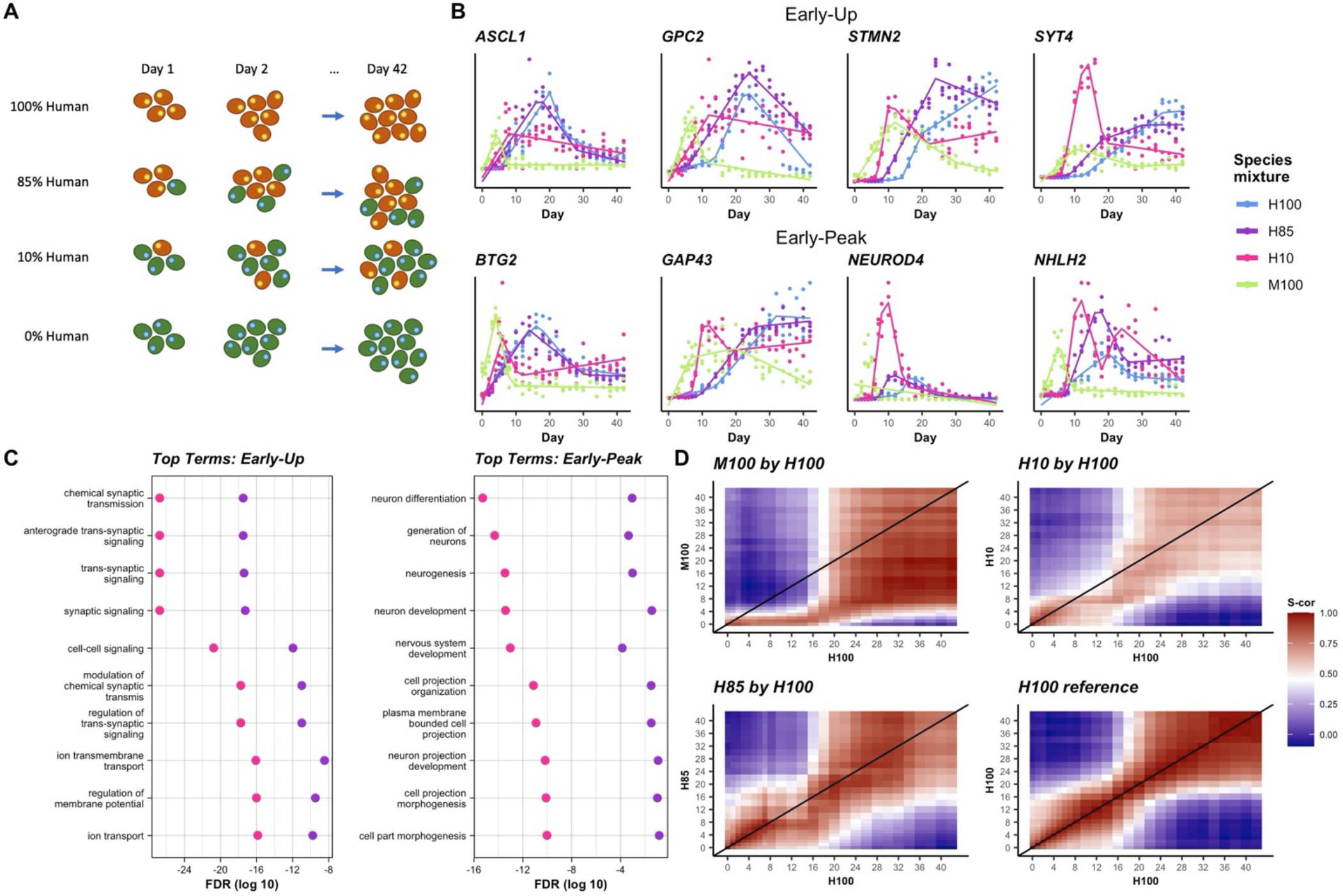
Variable mixing proportions show a dose response. (A) An additional, intermediate species mixing proportion is propagated and sequenced, denoted H85. (B) Expression plots of curated EU and EP genes with fitted trend lines (solid) for H100 (blue), H85 (purple), H10 (red), and M100 (green). Observed, normalized data are also plotted (dots). (C) Top 10 EU and EP GO terms from H10 showing relative significance of term enrichment for H10 and H85. (D) Correlation (Spearman) heat maps where regions of high correlation (red) below the diagonal indicate accelerated activity where later days in H100 are correlated with earlier days in the comparison mixture. Correlations are calculated on a subset of highly dynamic genes (see statistical methods).

Overall, expression profiles of a selection of key neuronal genes with either early up-regulation or early peaks in H85 samples were between H10 and H100 expression profiles (Fig. 6B). Overlaying these trends with the expression profiles of orthologous genes in the M100 sample reveals progressively later onsets of gene up-regulation/peaks with decreasing proportions of mouse cells among these genes (Fig. 6B).

To determine whether these results extended to the broader set of neuron-associated genes, we replicated the GO-term enrichment analysis in the H85 sample. Testing term enrichment on those genes which either up-regulated or peaked earlier in H85 relative to H100 resulted in a list of the most significant terms with the same patterns as in H10. However, comparing term significance levels between the top 10 most significant terms in the H10 analysis and their H85 counterparts shows that, while the H85 terms were still highly significant, they were less so than the H10 terms (Fig. 6C).

Pairwise correlations allowed us to further aggregate relative expression trends across terms. We took a subset of genes, targeting those with dynamic expression over time, and plotted correlations calculated between pairs of time points relative to H100 (Fig. 6D). Mouse orthologs demonstrate a visually significant acceleration with day 2 expression being highly correlated with H100 out to day 16. The H10 and H85 time courses both show visual acceleration with regions of high correlation below the diagonal, but with respectively lower magnitudes as the proportion of mouse cells decreases.

These analyses suggest that not only were the acceleration effects independent of simple culture parameters, but moreover that the effects were dose-dependent on the starting proportion of interspecies factors driving the acceleration.

### Human stem cells co-cultured with mouse cells correlated with *in vivo* human fetal neocortical samples earlier than human cells alone

We compared our data with human fetal sample references to assess if our *in vitro* acceleration is consistent with sample maturity *in utero*. The *Brain Span* database contains expression profiles from annotated brain regions across a range of developmental ages ^25–27^. We calculated correlations between our observed in vitro data and five tissue regions from the *Brain Span* database across weeks 8, 9, and 12 of development (see statistical methods for details). Across all time points and tissues, our mixed H10 and H85 samples increased correlation with the *Brain Span* reference earlier than the H100 control in a manner that was dose-dependent (Fig. 7).

**Figure 7:**
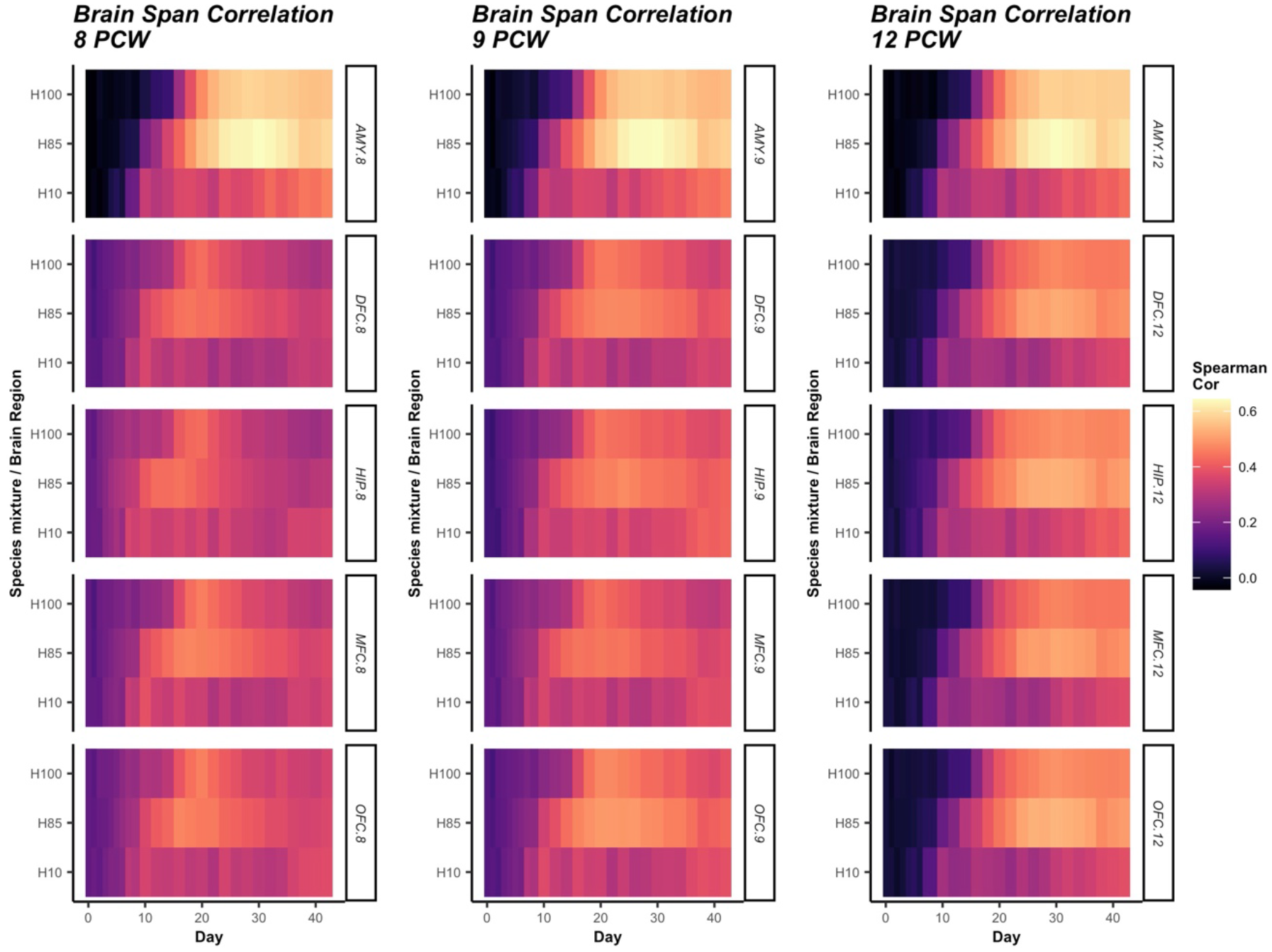
Correlation with Brain-Span regions further demonstrates dose response behaviors. Correlations (Spearman) between fitted trends and Brain-Span data are calculated at three Brain-Span time points and across the five brain regions represented at all time points. Calculations are performed on a subset of highly dynamic genes (see statistical methods).

These results were confirmed through a similar analysis of annotated brain tissue from the *Human Protein Atlas* ^28,29^ (S5 Fig.). As with the *Brain Span* data, higher proportions of mouse cells in the mixed samples resulted in earlier correlation with *in vivo* samples (S5 Fig.). These results are consistent with a genome-wide neural program that is activated earliest in M100, then significantly accelerated in H10, followed by moderately earlier in H85, and latest in H100 samples.

## DISCUSSION

In this study, we report for the first time multifaceted effects of interspecies mixing on the differentiation of hES cells. Through comprehensive RNA-seq time courses, we uncover that co-differentiation of hES cells intermixed with mEpiS cells was sufficient to accelerate components of neural gene regulatory programs, and identified genes with roles in neural lineage and regional identities that were both temporally and differentially expressed. We went on to demonstrate that the acceleration effect was dose-dependent on the starting ratio of interspecies cells (Fig. 6), and that the quickened expression patterns in chimeric samples correlated to *in vivo* tissue samples earlier in the differentiation time course than human samples alone (Fig. 7 and S5 Fig).

Accelerated neural developmental programs were indicated by earlier up-regulation of genes involved in neural migration and synaptic signaling (Fig. 3), as well as accelerated regulation of peak expression profiles of genes involved in neurogenesis (Fig. 4). Previously, we reported that the faster differentiation of mouse cells compared to human cells may be in part caused by increased speed of transcriptional upregulation of genes, indicated by steeper slopes in gene expression over time ^30^. Consistent with a mouse cell-induced acceleration of human cell neural differentiation, here we found that the slopes of peaked genes in human cells co-differentiated with mouse cells were also significantly increased in accelerated genes compared to control samples (S4B Fig.). However, non-peaking, mostly monotonic, genes whose upregulation began earlier showed lower slopes in chimeric samples, despite starting their upward trend significantly earlier and often continuing upwards for the duration of the time course (S4A Fig.). These results may suggest different functional roles of early-upregulated monotonic genes compared to genes with peak expression profiles. Indeed, genes with increased slopes and earlier peaks were significantly enriched in processes of generation of neurons and neuron cell projections, whereas earlier upregulated monotonic gene trends with lesser slopes were enriched in neuron and synaptic signaling events. Although we identify differences in gene expression profiles in our time course in this report, the functional maturity of resulting neurons in control versus chimeric co-differentiation conditions remains to be determined.

The mechanisms responsible for both the developmental clock and how interspecies co-culture may affect the differentiation speed of another species remain unknown. Recently, *in vitro* models of mouse and human segmentation clocks with species-specific timing has been reported ^31–34^. Although the driver of a universal developmental clock for all tissues is unknown, it has been speculated that, in the case of the segmentation clock, differences in *HES7* gene expression and protein degradation rates control oscillation frequencies that drive the rate of somitogenesis. It has been proposed that metabolic and biochemical reaction rates in cells of different species might modulate developmental rates ^35^. Here, we show that cell-cell signaling alone is sufficient to affect the developmental clock.

Previously, several studies suggested that the intrinsic species-specific developmental timer was faithfully retained under various conditions. First, the intrinsic developmental clock seemed independent of culture method as both 2D culture and 3D organoid systems exhibited similarly robust retention of developmental time^9,10,36–38^. Second, several chimeric transplant/implantation studies also suggested the retention of developmental time of the species of origin rather than the host. For example, earlier our group reported that the developmental rate of hES cell-generated teratomas strictly retained human developmental time despite being grown in a mouse host ^4^. Multiple other labs have also reported that transplantation of hES cell-derived neural progenitor cells into mouse brains was insufficient to accelerate the timing of human neural maturation ^6,7,10^. Previous *in vitro* interspecies co-culture of stem cell-derived neural cells from different primate species failed to demonstrate deviation from intrinsic developmental properties in one study, however the mixed progenitor cells in this instance were already well differentiated towards neural cell types ^39^. While these studies revealed that non-embryonic interspecies conditions were insufficient to alter developmental time, in this study we demonstrate that factors actively driving an embryonic developmental program from pluripotency, rather than a mature host environment, can be sufficient to affect components of the developmental clock of cells from another species.

The ability of stem cells of different species to resolve conflicting developmental speeds has significant implications in the development of chimeric embryos for human organ formation ^40^. With a widespread shortage of immunologically-matched organs for patients in need of organ transplants, the ability to grow transplantable human organs through human stem cell chimeric contributions to embryos remains an interesting potential therapeutic approach ^41,42^. However, many barriers remain, including poor human chimeric contributions, possibly in part due to the vastly different developmental rates between neighboring cells of different species ^11,40,43^. In this study, we demonstrate that it is possible for mouse cells to influence developmental rates and outcomes of neighboring human cells.

Previous reports of successful human cell contributions to chimeric mammalian embryos ^41,44,45^, including a recent report of the highest contribution (4%) of human cells in mouse-human chimeric embryos ^46^, could imply that human pluripotent stem cells may be induced to accelerate their developmental rate to match that of their embryonic host species. However, maturation rates of human cells in interspecies chimeras have not been well characterized. Our comprehensive time course results in this study indicate that human developmental time could be accelerated by co-differentiating cells within chimeric embryos, although collateral impacts in cell lineage outcomes may occur. In the case of neural differentiation in this study, we did find genes involved in dorsal forebrain development, for example, that were temporally downregulated in interspecies samples while genes involved in ventral midbrain development were upregulated, likely, at least in part, due to an earlier and increased exposure to SHH (Figs. 3–5) ^47–49^. Importantly, mouse and human brains do not share identical brain physiologies, cell type compositions, nor brain region proportions ^50,51^, so it is perhaps not surprising that altered cell fate choices are made when cells are exposed to signals intended to created divergent outcomes. Thus, it will be important to monitor cell outcomes in chimeric embryos for human organ growth to verify that cell type contributions and organ functions are not affected.

Although the protocol described here will not have clinical applications due to the xenotropic nature of the conditions, it does suggest that the human developmental clock can be accelerated. Although the specific factors involved and clock mechanism itself remain to be dissected, this proof-of-concept report provides evidence that the species-specific developmental clock may be amenable to acceleration for clinically-relevant benefit.

## Materials & Methods

### Cell culture

Human ES and mEpiS cells were cultured and passaged as previously reported^4^. Briefly, H9 cells were cultured in E8 Medium (Thermo Fisher Scientific, USA) on Matrigel-coated plates and split every 2-3 days with EDTA. To easily identify human from mouse cells, H9 cells were electroporated with a selectable PiggyBAC-inserted plasmid expressing nuclear-localized H2B-mCherry driven by the EF1α promoter, and clonally expanded. EGFP-expressing mEpiS cells derived from C57BL/6-Tg(CAG-EGFP)1Osb/J (JAX Stock No. 003291) mice and cultured as previously described^4,19,21^. Cell were maintained on low passage MEFs and cultured in DMEM/F12 medium (Thermo Fisher Scientific, USA) supplemented with 20% Knockout serum replacement (Thermo Fisher Scientific, USA), 0,18 mM B-mercaptoethanol (Sigma, USA), 1Xnon-essential amino acids (Thermo Fisher Scientific, USA), 2 mM L-glutamine (Sigma, USA), 7.5 ng/mL activin A (R&D Systems, USA), and 5ng/mL bFGF (R&D Systems, USA). Cells were passaged by adding TrypLE (Thermo Fisher Scientific, USA) and seeding onto fresh MEFs with 10 μM Y27632 ROCK inhibitor overnight to increase cell survival (Tocris Bioscience, UK).

### Neural induction and sampling for RNA-seq

At day 0 of time courses, H9-H2BmCherry and EGFP-mEpiS cells were washed with PBS (Thermo Fisher Scientific, USA), treated with TrypLE (Thermo Fisher Scientific, USA) for singularization, and resuspended in a simple neural differentiation medium consisting of DF3S (DMEM/F-12, L-ascorbic acid-2-phosphate magnesium (64 mg/L), sodium selenium (14 μg/L), and NaHCO3 (543 mg/L), Thermo Fisher Scientific, USA), 1XN2 supplement (Thermo Fisher Scientific, USA), 1XB27 supplement (Thermo Fisher Scientific, USA), and 100ng/mL of mNoggin (R&D Systems, USA). To aid cell survival, 10 μM Y27632 ROCK inhibitor (Tocris Bioscience, UK) was added on day 0, and cells were mixed at the indicate mouse-human ratios and seeded into Matrigel-coated 12-well plates at 2.5X10^5^ cells/well in triplicate. Media in all wells was replaced with fresh neural differentiation media (without ROCK inhibitor) every day for the 42 days of differentiation. When cells become over-confluent cells were split 1:3 or 1:6 by EDTA-treatment to avoid disrupting cell-cell interactions.

### Flow Cytometry, Microscopy, and Time Lapse Imaging

Human-mouse cell ratios were established by monitoring red and green fluorescence, respectively, by flow cytometry. Cells were treated with 350μL TryPLE, spun down, and resuspended in 400 μL FACS buffer (PBS + 5% Bovine Serum Albumin). Cells were analyzed on a BD FACSCanto II and analyzed using FlowJo 9.3 software (Becton Dickinson & Company, USA).

All time-lapse microscopy was acquired on a BioStation CT automated imaging system (Nikon Instruments, Japan). Samples from all conditions were imaged at least every other day using phase-contract and fluorescence microscopy. For time-lapse movies, cells were acquired with a 10X magnifying objective every 30 minutes for the first 6 days of differentiation in phase-contrast and green and red fluorescence channels. Overlaid movies were compiled with CL-Quant software (DRVision, USA).

### Sample processing and RNA-seq pipeline

For RNA sample collection, samples were washed with 1XPBS (Thermo Fisher Scientific, USA) and lysed in 700 μL RLT-PLUS buffer (Qiagen, USA), and stored at −80C until further processing. Total RNA was then purified from 350 μL RLT-Plus Buffer using RNeasy Plus 96 and Micro Kits (Qiagen, Netherlands) and quantitated with the Quant-iT RNA Assay Kit (Thermofisher, USA). RNA was diluted to one hundred nanograms for input. The Ligation-Mediated Sequencing (LM-Seq) protocol was used to prepare and index all cDNA libraries (Hou et al 2015). Final cDNA libraries were quantitated with the Quant-iT PicoGreen Assay Kit (Thermofisher, USA). Twenty-five to forty-eight uniquely indexed samples were pooled per lane on an Illumina HiSeq 2500 with a single 51 base pair read and a 10 base pair index read.

A joint hg19/mm10 transcriptome reference was built by appending hg19 or mm10 respectively to the chromosome sequences and gene symbols. Tagging the gene symbols with the ID of the reference genome ensured easy decomposition of the resulting expression estimates into mouse and human subsets of species-specific gene expression. Mitochondrial genes were removed prior to further downstream analysis or normalization due to their inconsistent abundance across samples.

The sequencer outputs were processed using Illumina’s CASAVA-1.8.2 base calling software. Sequences were filtered and trimmed to remove low quality reads, adapters, and other sequencing artifacts. The remaining reads were aligned to the joint transcriptome using RSEM version 1.2.3 with bowtie-0.12.9 for the alignment step. After ensuring accurate mapping to the human/mouse subset of the transcriptome (see below for details), identified by the respective hg19 and mm10 tags on the gene symbol, the human and mouse subsets of expected counts were separated for individual analysis.

### Mixed species sample quality control

To assess the quality of alignment to the combined human-mouse transcriptome, misalignment rates were quantified in the H100 (pure human) and M100 (pure mouse) samples. In these cases, transcripts which align to the mouse and human subset of the transcriptome respectively represent errors of misalignment. Typical misalignment rates across samples appeared to be well controlled as the majority of H100 samples aligned less than 0.5% of transcripts to mouse genes (median ^~^0.35%, third quartile ^~^0.37%). The majority of M100 samples similarly aligned less than 1.5% of transcripts to human genes (median ^~^0.53%, third quartile ^~^1.42%) (S2 Fig.).

A few samples (^~^5%) exhibited high misalignment rates (>5%). For this reason, samples with unusually low sequencing depth were removed. The filtering criteria considered log10 transformed sequencing depth (within sample sum of total expression) and removed samples with depth below the median minus 1.5 times the IQR. This procedure removed the majority of individual samples in H100 and M100 with high alignment error rates. Therefore, misalignment is believed to be primarily a function of, or at least well predicted by, low sequencing depth (S2 Fig.).

A second filter was implemented to remove samples with expression profiles significantly different from biological replicates of the same time point and temporally neighboring samples. Normalized data (see below for details) from the top 1000 highest variance genes across samples within each mixture was reduced to 10 principal components. This number roughly accounts for the majority of temporal variability based on the variance explained by each component. Loadings for each component were expected to follow a smooth curve in time, following the portion of the developmental trajectory defined by the principal component. For this reason, loadings were fitted with a 4^th^ degree spline regressed against time. Studentized residuals were tested for being significantly different than the regression curve. A sample level p-value was derived by testing against the null distribution that the maximum residual across the 10 components (in absolute value) was t-distributed. The method of Benjamini and Hochberg^52^ was used to provide adjusted p-values. A backward elimination and forward selection procedure was then applied. Specifically, the sample with the smallest adjusted p-value below 1e-05 was removed and the process repeated until no samples had an adjusted p-value below 1e-05 (if a sample is the last remaining observation from a particular time point, it was not considered for removal regardless of its adjusted p-value). Samples were then added back in one-at-a-time in the order of removal. Any with adjusted p-values above 1e-05 were retained for further analysis, and otherwise were rejected permanently. The filtered dataset was renormalized prior to analysis.

Empirically, this procedure was shown to remove several remaining high-error samples from M100 without removing high sequencing depth samples across species mixture groups (S2 Fig.).

### Normalization of mixed species samples

We used a modified application of the scran^53^ method for normalization of the expected count data. Human and mouse aligned transcripts were normalized separately, and so relative levels of normalized expression were not directly comparable between species. Consider the human mixtures (H10, H85, or H100); mouse mixtures were normalized identically. When biological replicates existed for a time point, scran was first applied to normalize these samples. Average normalized expression of biological replicates was then normalized, again via scran, across both time points and mixtures.

### Segmented regression and gene-trend classification

The dynamics of gene expression through time were defined by a segmented regression implemented using the Trendy^22^ package. Trendy automatically selects the optimal number of segments (up to a maximum of 5 in this application) and requires that each segment contain a minimum number of samples (5 in this application). Additionally, an automatic significance test on segment slopes classifies segments as increasing, decreasing, or flat. As the test is itself somewhat conservative, we used a significance threshold of 0.1 (default) to determine these slope classifications. Trendy was then applied to all genes for which the 80% quantile of normalized expression is above 20 for at least one mixture.

Following regression, the segment trend classifications were used to define sets of genes by patterns of behavior relative to a reference dataset (H100 in the majority of the published analysis). Genes were classified into subsets of accelerated or differentially expressed (DE) relative to the reference dataset according to the following criteria:

1. Accelerated by Early Up (EU):
  a. Both the test gene and the reference gene contain an increasing segment which is not preceded by a decreasing segment. If multiple such segments exist, only the first is considered.
  b. The increasing segment in the test gene must start at least 2 days before the increasing segment in the reference gene.
  c. The slope of the increasing segment in the test gene must be at least 5 times the slope of the (non-increasing) reference segment which contains the start time of the test increasing segment (typically the segment just prior to the increasing reference segment). This filter removes genes for which the reference segment containing the start time is labeled as flat by Trendy (slope is not significantly different from 0), but is fitted with an up-trending slope. This can happen in instances where the reference segment is short and so does not contain enough sample points for the up-trend to be labeled as significant.
2. Accelerated by Early Peak (EP):
  a. Both the test gene and the reference gene contain a peak defined by an increasing segment followed by a flat or decreasing segment. The peak itself is defined by the time of the breakpoint between these two segments.
  b. The peak in the test gene must be at least 2 days before the peak in the reference gene.
3. DE Up:
  a. The maximum fitted value of the test gene plus 1 must be at least 3 times the maximum fitted value of the reference gene plus 1. The inclusion of the plus 1 bias to each side prevents very lowly expressing genes from appearing DE due to small differences in fitted values which are only multiplicatively large due to the low overall expression.

Genes in H10 or H85 matching these acceleration/up-regulation criteria were denoted as “Early” or “Up” respectively.

We also ran this classification denoting H100 as the test datasets. When genes matched the criteria in this case, we denoted the corresponding gene in the reference dataset, H10 or H85, “Late” or “Down” according to the specific criteria met.

### Gene set enrichment

Accelerated and DE gene sets were further characterized through testing for GO term enrichment. The topGO^54^ package and org.Hs.eg.db^55^ dataset were used to perform enrichment testing on GO terms belonging to the biological processes (BP) ontology. The set of all genes on which Trendy segmented regression was run was used as the background set (see above for subset definition). Significant p-values were then FDR corrected^52^ prior to analysis.

### Correlation analysis

Expression similarity across time points, species mixtures, and external reference datasets was assessed through gene expression correlations. To ensure that computed correlations were representative of the temporal gene dynamics being studied, correlations were computed on only a subset of genes. Highly dynamic genes were subset from all Trendy-fit genes by calculating the coefficient of variation of fitted values. The highest CV across species mixtures was then retained as a measure of each gene’s level of temporal dynamics, and the top 1000 most dynamic (highest CV) genes were subset for analysis.

Relative acceleration of species-mixtures was computed as the correlation matrix (spearman type) between time points where within-day technical replicates were averaged together to obtain a single day expression value.

Relative acceleration in combination with brain-region similarity on the species-mixture data was then separately validated/assessed through similar calculation of correlations between the species-mixture data and two outside datasets: the BrainSpan atlas of the developing human brain^26,27^ and the Human protein atlas^28,29^.

### R package versions

All calculations were performed using R^56^ (v3.6.2) and major packages: Trendy^22^ (v1.6.4), scran^53^ (v1.12.1), topGO^54^ (v2.36.0), org.Hs.eg.db^55^ (v3.8.2), ggplot2^57^ (v3.3.0).

## List of Abbreviations

(RNA-seq): RNA-sequencing
(hES) cells: human embryonic stem
(mEpiS) cells: mouse epiblast stem
(LM-Seq): Ligation-Mediated Sequencing
(GO): Gene Ontology

## Declarations

### Ethics approval and consent to participate

All experiments described in this study were approved by the ethics committee with IRB Approval Number: SC-2015-0010. The H1 hES cells are registered in the NIH Human Embryonic Stem Cell Registry with the Approval Number NIHhESC-10-0043.

### Consent for publication

Not applicable.

### Data Availability

The RNA-seq datasets supporting the conclusions of this article are available (pending publication) in the Gene Expression Omnibus repository, GSE157354. The code for reproducible analyses and generation of figures and tables is available (pending publication) at https://github.com/JBrownBiostat/ChimericDevelopment.

## Supporting information

**S1 File. Summaries of expression characteristics for genes classified as exhibiting differential timing or expression in H10.**

**S2 File. Summaries of expression characteristics for genes classified as exhibiting differential timing or expression in H85.**

**S1 Movie:** H9-H2BmCherry (red) cells mixed 1:10 with EGFP+ mouse EpiS cells (green) were seeded in neural differentiation medium and imaged every 30 mins for the first 6 days of differentiation using a Nikon BiostationCT imaging system. Condensation was noted during the first images capture after media replacement approximately every 24 hours. Overlaid channels of microscopy images were compiled into the movie with CL-Quant software (DRVision, USA).

**S1 Fig. Quality control filtering removes samples with uncharacteristically low sequencing depth.** (Top) Observed per-sample misalignment rates for pure human/pure mouse mixtures. (Middle/Bottom) Observed log10 total sequencing depth summed across sequences aligned to either human or mouse. Most samples removed from analysis (blue) are below the depth filtering threshold (dashed line) (see statistical methods). Otherwise, the M100 results suggest that the higher-depth removed samples are those with higher rates of misalignment (top/middle, right column).

**S2 Fig. Seeded human cell proportions increase over time.** (A) Observed percent of human cells in H10 mixture out to 16 days. (B) FACS intensities used to compute relative proportions of human and mouse cells in H10 mixture.

**S3 Fig. Selected gene expression plots show characteristic differences between H100, H10, and M100.** (A) EU classified fitted trend lines (solid) are plotted for selected genes with overlaid normalized observed data (points). (B) Similar results are shown for selected EP classified genes.

**S4 Fig. Up-trends show defining shifts in H10 among EU and EP genes.** (A) Slope ratio (ratio of H10 up-trend slope over H100 up-trend slope) densities are plotted (left) on the log scale for top enriched GO terms. KS testing shows a significant left-shift corresponding to significantly reduced slopes in H10 among these genes. Densities of the duration of up-trends (right) show significantly longer (KS test) trends for H10 (red) than H100 (blue). (B) Similar results for EP genes show significant increases in slope in H10 with reduced duration of up-trend.

**S5 Fig. Correlation with Human Protein Atlas (HPA) data further demonstrates dose response behaviors.** Correlations (Spearman) between fitted trends HPA data are calculated across the thirteen HPA regions. Calculations are performed on a subset of highly dynamic genes (see statistical methods).

## Acknowledgements

The authors would like to thank John Maufort for editorial assistance and critical reading of this manuscript.

## Author Contributions

Conceptualization: Jared Brown, Christopher Barry, Matthew T. Schmitz, James A. Thomson, Christina Kendziorski.

Data curation: Jared Brown, Matthew T. Schmitz, John Steill, Scott Swanson.

Formal analysis: Jared Brown, Christopher Barry, Matthew T. Schmitz, Michael Schwartz, and Christina Kendziorski.

Funding acquisition: James A. Thomson and Christina Kendziorski.

Investigation: Christopher Barry, Cara Argus, Jennifer M. Bolin, Amy Van Aartsen.

Methodology: Jared Brown, Christopher Barry, Matthew T. Schmitz, Scott Swanson, Michael Schwartz, James A. Thomson, Christina Kendziorski.

Project administration: Christopher Barry and James A. Thomson.

Resources: Ron Stewart, James A. Thomson, and Christina Kendziorski.

Software: Jared Brown, John Steill, Scott Swanson, and Christina Kenzdiorski.

Supervision: Christopher Barry, James A. Thomson, and Christina Kendziorski.

Validation: Jared Brown, Christopher Barry, John Steill, Scott Swanson, Ron Stewart, and Christina Kendziorski.

Visualization: Jared Brown, Christopher Barry, and Matthew T. Schmitz.

Writing – original draft: Jared Brown and Christopher Barry.

Writing – review & editing: Jared Brown, Christopher Barry, Matthew T. Schmitz, Jennifer M. Bolin, John Steill, Scott Swanson, Ron Stewart, James A. Thomson, and Christina Kendziorski.

