## Supplemental figures for "Interspecies Chimeric Conditions Affect the Developmental Rate of Human Pluripotent Stem Cells"

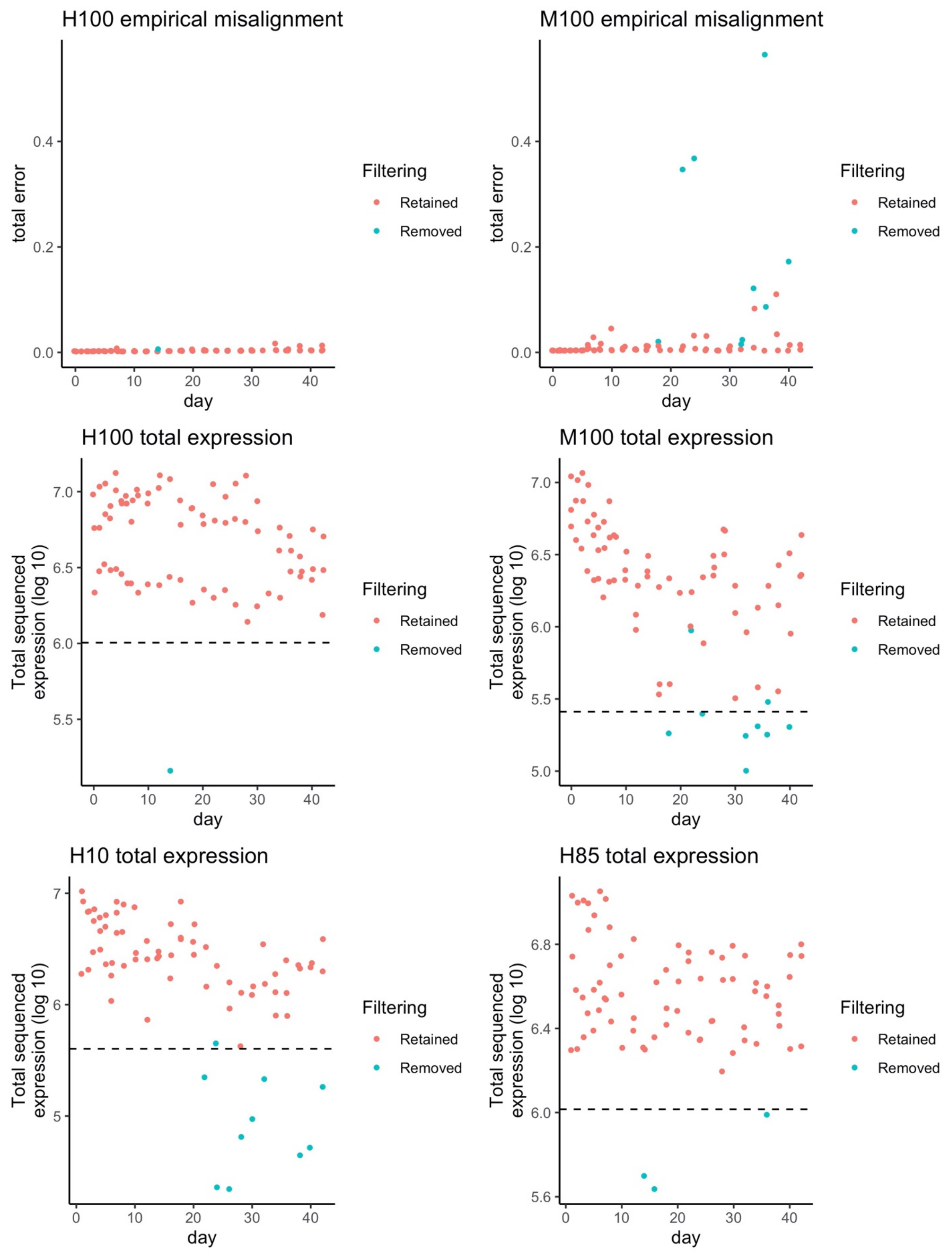


*Supplemental Figure 1: Quality control filtering removes samples with uncharacteristically low sequencing depth. (Top) Observed per-sample misalignment rates for pure human/pure mouse mixtures. (Middle/Bottom) Observed log10 total sequencing depth summed across sequences aligned to either human or mouse. Most samples removed from analysis (blue) are below the depth filtering threshold (dashed line) (see statistical methods). Otherwise, the M100 results suggest that the higher-depth removed samples are those with higher rates of misalignment (top/middle, right column).*

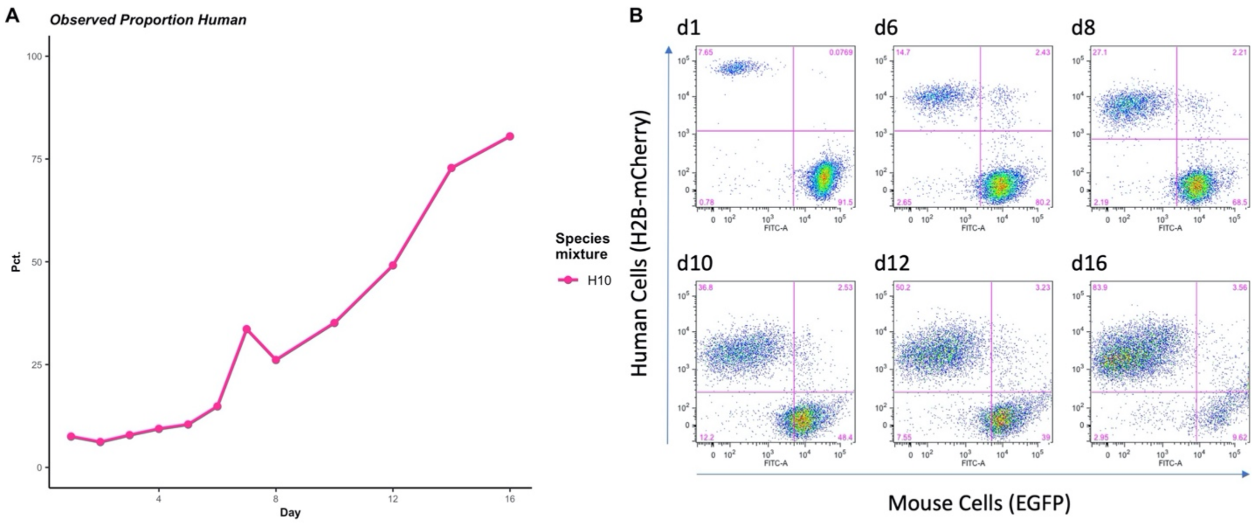


*Supplemental Figure 2: Seeded human cell proportions increase over time. (A) Observed percent of human cells in H10 mixture out to 16 days. (B) FACS intensities used to compute relative proportions of human and mouse cells in H10 mixture.*

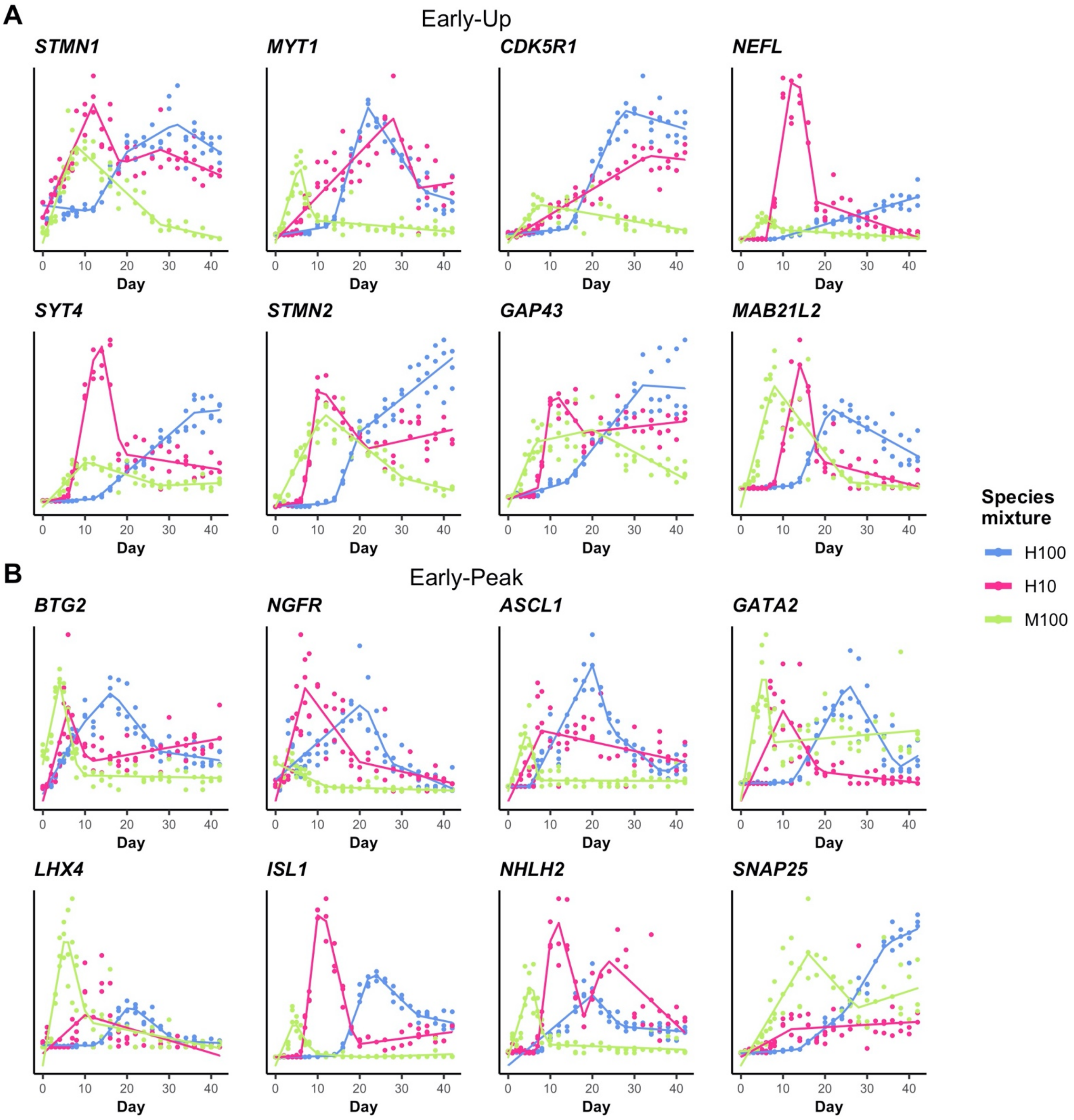


*Supplemental Figure 3: Selected gene expression plots show characteristic differences between H100, H10, and M100. (A) EU classified fitted trend lines (solid) are plotted for selected genes with overlaid normalized observed data (points). (B) Similar results are shown for selected EP classified genes.*

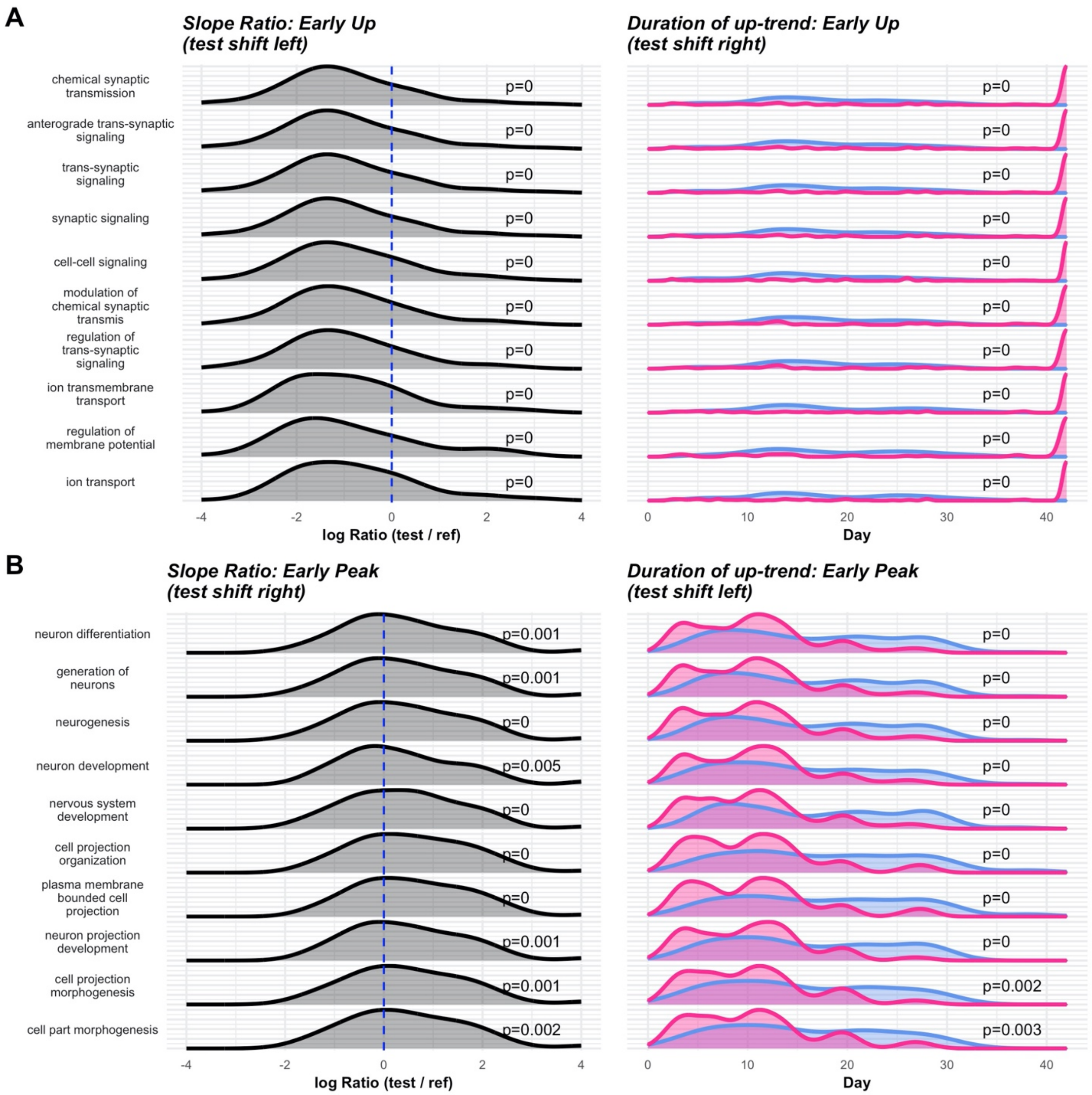


*Supplemental Figure 4: Up-trends show defining shifts in H10 among EU and EP genes. (A) Slope ratio (ratio of H10 up-trend slope over H100 up-trend slope) densities are plotted (left) on the log scale for top enriched GO terms. KS testing shows a significant left-shift corresponding to significantly reduced slopes in H10 among these genes. Densities of the duration of up-trends (right) show significantly longer (KS test) trends for H10 (red) than H100 (blue). (B) Similar results for EP genes show significant increases in slope in H10 with reduced duration of up-trend.*

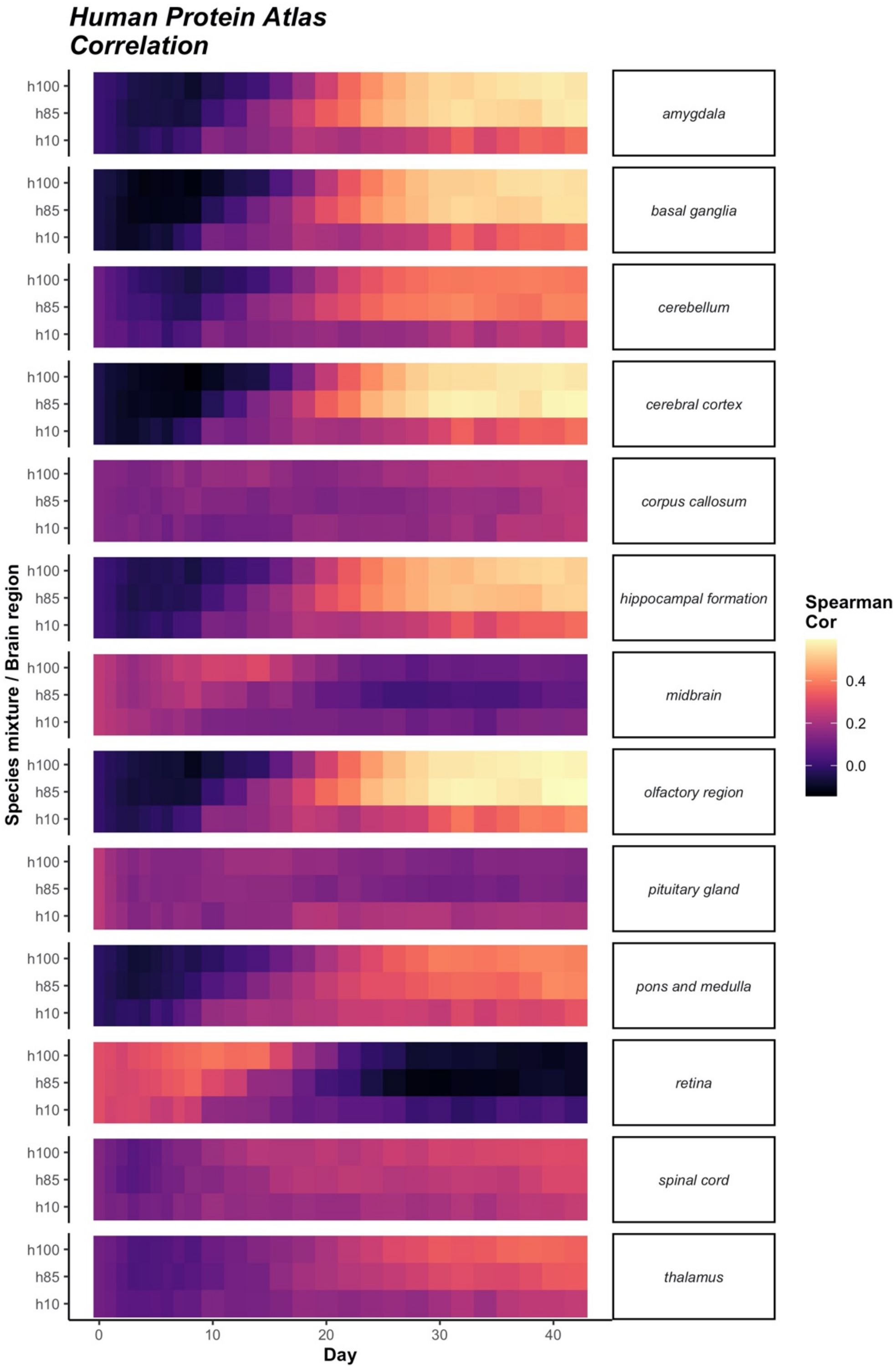


*Supplemental Figure 5: Correlation with Human Protein Atlas (HPA) data further demonstrates dose response behaviors. Correlations (Spearman) between fitted trends HPA data are calculated across the thirteen HPA regions. Calculations are performed on a subset of highly dynamic genes (see statistical methods).*
